# Tissue-Specific Prevalence and Clonal Architecture of *BRCA1/2* LOH-Inducing Chromosomal Aneuploidy

**DOI:** 10.64898/2026.04.17.718766

**Authors:** Xinfeng Wang, Saumya D. Sisoudiya, Mahad Bihie, Yves Greatti, Julián Grandvallet Contreras, Tomi W. Jun, Smruthy Sivakumar, Kuan-lin Huang

## Abstract

Germline pathogenic variants in BRCA1 and BRCA2 confer disproportionately elevated cancer risks in breast and ovarian tissues, yet the basis for this tissue specificity remains incompletely understood. Here, we integrate bulk-tumor aneuploidy analysis across 340,824 cancer cases from three independent cohorts (TCGA, ICGC PCAWG, and FoundationCore) with single-cell whole-genome sequencing from two independent studies to investigate whether tissue-specific patterns of chromosomal deletion contribute to this phenomenon. We find that breast and ovarian cancers are consistently enriched for deletions of chromosome arms 17q and 13q—harboring the *BRCA1* and *BRCA2* genes, respectively—relative to other solid tumor types, and that mutational timing analysis independently places these deletions among the earliest somatic events in these cancers. Phylogenetic reconstruction of single-cell data reveals that in pre-malignant breast tissue from germline *BRCA1/2* carriers, chr17q and chr13q deletions appear as localized subclonal events within small clades against a largely diploid background. In established malignancies, these same deletions are found within dominant clonal lineages accompanied by widespread genomic instability—consistent with clonal sweeps originating from early deletion events. These findings suggest that breast and ovarian cellular environments confer a selective advantage for chr17q and chr13q deletions, providing a mechanism that may contribute to the tissue-specific cancer risk observed in gBRCA1/2 carriers.

## MAIN TEXT

Pathogenic germline *BRCA1/2* variants^1,2^ (abbreviated *gBRCA1/2* variants below) are inherited and present across cells within a carrier individual, yet their associated cancer risk displays a striking predominance towards breast and ovary tissues^3,4^. The standardized incidence ratios (SIRs) for breast or ovarian cancer in women carrying *gBRCA1/2* were at least 12.9 across ages^5^. The cumulative breast cancer risk by age 80 was 72% for *gBRCA1* carriers and 69% for *gBRCA2* carriers^5^, and the cumulative ovarian cancer risk by age 80 was 44% for *gBRCA1* carriers and 17% for *gBRCA2* carriers. In comparison, the relative risks (RRs) of *gBRCA1/2* carriers, compared to non-carriers, were less than 5 for any other single cancer type^6–9^.

The reason underlying this tissue-specificity for *BRCA*-associated cancer risk remains unresolved^10^. Previous studies examining tissue-specific splicing, gene and protein expression^11,12^, as well as tissue-microenvironment factors including transcriptional programs, hormones, or metabolites^10^ have yielded inconclusive results. Estrogen, commonly found in breast and ovarian tissues, may induce double-strand breaks (DSBs) and could exacerbate *gBRCA1/2* deficiency^13,14^, potentially increasing cancer risk. However, estrogen levels are also high in selected tissues like the uterus, where *gBRCA1/2*-associated cancer risk does not match that of breast and ovary. Additionally, other tissues are also frequently exposed to DSB-associated carcinogenic agents^15^ or present with DSB signatures^16^, but do not show a comparable cancer risk.

Based on the two-hit hypothesis formulated by Alfred G. Knudson^17^, *gBRCA1/2*-mediated oncogenesis is thought to originate from a cell where the *gBRCA1/2* variant underwent loss of heterozygosity (LOH) of the wildtype *BRCA1/2* allele, resulting in homologous recombination (HR) DNA repair deficiencies and genomic instability^18,19^. Supporting this theory, LOH of *BRCA1* and *BRCA2* variants was observed in 80 out of 90 germline or somatic *BRCA1/2* breast cancer cases^20^, likely resulting in loss of HR function^21^. While LOH can occur through multiple mechanisms—including focal deletion, copy-neutral LOH (cn-LOH), and arm-level or whole-chromosome deletion—arm-level chromosomal aneuploidy is among the most common routes and exhibits distinct tissue-specific patterns across cancer types^32^.

We hypothesized that the tissue-specific patterns of chromosomal aneuploidy, specifically deletions affecting the chromosome arms harboring *BRCA1* (17q) and *BRCA2* (13q), may represent a tissue-dependent mechanism that contributes to the higher rate of second-hit LOH events in breast and ovarian tissues. To test this hypothesis, we integrated large-scale bulk-tumor cohort analyses with single-cell phylogenomic reconstruction, the latter providing direct evidence of the clonal architecture and evolutionary timing of these deletions that has not been previously examined in this context.

## RESULTS

### Aneuploidy affecting *BRCA1/2*-located chromosome arms, chr17q and chr13q

*BRCA1* and *BRCA2* genes are located on chromosome arms 17q and 13q (abbreviated chr17q and chr13q), respectively. We note that these arms also harbor other important tumor suppressor genes (TSGs)—including RB1 (13q14.2), NF1 (17q11.2), and CDK12 (17q12)—whose loss may independently contribute to selective advantage for these deletions. TP53, located on 17p, may also be co-deleted in cases of whole-chromosome 17 loss. To characterize tumor aneuploidy affecting the *BRCA1/2*-located chromosome arms, we analyzed and compared the prevalences of chr17q and chr13q deletions across tumor types in three major cancer genomics cohorts totaling 340,824 cases: The Cancer Genome Atlas (TCGA, N = 10,522), the International Cancer Genome Consortium Pan-Cancer Analysis of Whole Genomes datasets (ICGC PCAWG, N = 1,729), and FoundationCore® database from Foundation Medicine Inc. (FCore, N = 328,573) (**Supplementary Table 1A-C** with detailed cancer type abbreviations). In TCGA and ICGC, this analyzed sample size reflected only one primary tumor per patient that had no overlapping samples between the two datasets. The FCore cohort also included only one tumor sample per patient.

In TCGA primary tumors, ovarian and breast cancer cases showed significantly higher prevalences of chr17q deletion (70.2% and 19.0%, respectively, **Figure 1A, Supplementary Table 2A**). We devised a permutation test to assess whether the observed prevalences of chr17q arm-level deletion were higher than a random-chance cancer type within TCGA cohort (**Methods**), these two cancer types, along with ACC and KICH, were the only ones that showed significantly higher prevalence (FDR < 1e-4). For chr13q deletion, ovarian and breast cancer also demonstrated significantly higher prevalence (56.9% and 40.6%, Permutation test FDR < 1e-4, **Figure 1A**). Multiple other cancer types showing significantly higher chr13q deletion prevalences, including LUSC, MESO, LUAD, and HNSC, were classified as cancer types with high environmental influence^22^, which may drive the higher rates of aneuploidy seen in these cancer types. In the ICGC PCAWG dataset that excluded TCGA samples, primary tumors of ovarian and breast cancer cases showed significantly higher prevalences of chr17q deletion (37.7% and 24.8%, respectively, Permutation test FDR < 1e-4 and FDR = 0.0045, respectively, **Figure 1B, Supplementary Table 2B**). For chr13q deletion, ovarian and breast cancer also demonstrated relatively higher prevalences (26.2% and 30.6%, respectively, FDR = 0.0045 and FDR < 1e-4, respectively, **Figure 1B**).

**Figure 1.**
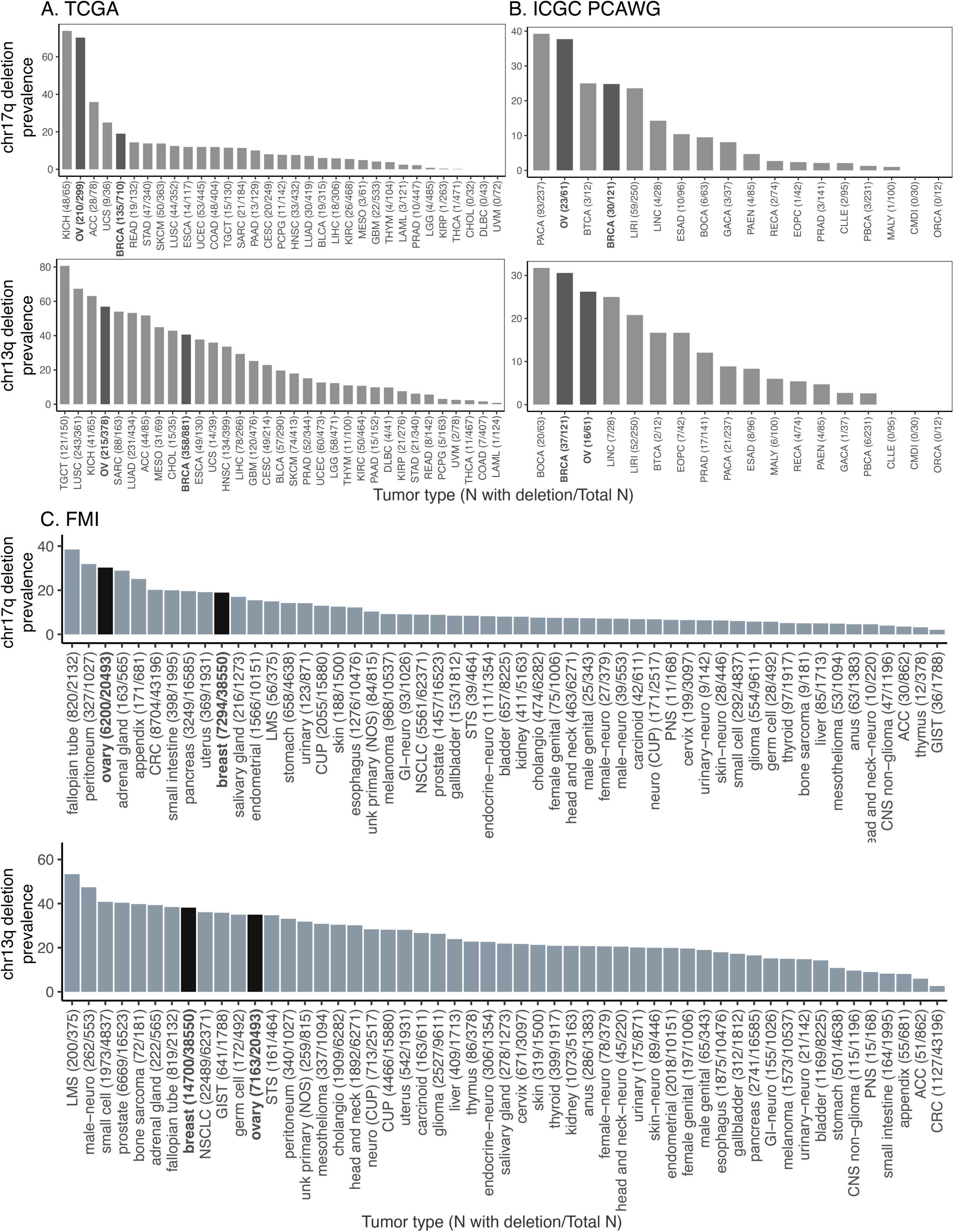
The prevalence of deletions in chromosome arms 17q and 13q across tumor types in (A) TCGA, (B) ICGC PCAWG, and (C) FCore cohorts. The x-axis lists the tumor types along with the number of samples with deletions out of the total number of samples analyzed, e.g., “CANCER TYPE (#N affected/#Total N)”. The y-axis indicates the percentage of samples with the specified deletion for each tumor type. Tumor types with a higher prevalence of deletions are shown on the left, decreasing towards the right.

A fraction of tumors undergo whole-genome doublings (WGD); in such circumstances, a heterozygous deletion of chr13q/17q deletion may not confer LOH to the *BRCA1/2* variants. To rule out these cases, we removed samples with WGD and recalculated the prevalences of chr17q and chr13q deletions in TCGA and ICGC. In TCGA, non-WGD ovarian and breast cancer still showed significantly higher prevalences of chr17q deletion (71.4% and 11.2%, respectively, Permutation test FDR < 1e-4, **Figure S1A, Supplementary Table 2A**). For chr13q deletion, ovarian and breast cancer continued to show relatively higher prevalences, after excluding WGD cases (51.0% and 24.9%, respectively, Permutation test FDR < 1e-4, **Figure S1A**). In the ICGC PCAWG dataset, ovarian and breast cancer showed higher prevalences of chr17q deletion (82.6% and 33.8%, respectively, Permutation test FDR < 1e-4 and FDR = 0.0017, respectively, **Figure S1B, Supplementary Table 2B**). Similarly, ovarian and breast cancer showed higher prevalences of chr13q deletion (60.9% and 27.9%, Permutation test FDR < 1e-4 and FDR = 0.00067, respectively, **Figure S1B**). These results show that the higher prevalences of chr17q and chr13q deletions observed in ovarian and breast cancer were not affected by WGD status.

The FCore cohort analyzed herein comprised 328,573 patients with solid tumors who received tissue biopsy-based comprehensive genomic profiling (CGP) using FoundationOne® (n = 87,807) or FoundationOne®CDx (n = 240,766) during routine clinical care. The samples received for profiling may comprise primary, metastatic or lymph node biopsies. Overall, 25.9% of patients had a loss of chr13q and 14.1% had a loss of chr17q. Among the 53 tumor types with at least 100 total samples, 19 tumor types had a >10% prevalence of chr17q loss (**Figure 1C, Supplementary Table 2C**). The prevalence of chr17q loss in ovarian and breast cancer was 30.3% (6,200/20,493) and 18.9% (7,294/38,550), respectively. Given that this cohort contains a more comprehensive representation of tumor types with sufficient sample size, we devised a more stringent permutation test (**Methods**) to test the null hypothesis that ovarian or breast cancer ranks higher in prevalence across solid cancer types than in the permuted cohorts. These cancer types showed significantly higher prevalence of chr17q loss compared to other cancer types in this dataset (ranked at 3/53 and 10/53, p < 1e-4 and p = 0.0029, respectively, Permutation test). The only two cancer types that showed significantly higher prevalence of chr17q loss above ovarian cancer are fallopian tube and peritoneum cancers, which are adjacent and may share common origins with ovarian cancer. For chr13q loss, 47 of the 53 tumor types with at least 100 total cases showed a prevalence greater than 10% (**Figure 1C, Supplementary Table 2C**). The prevalence of chr13q loss in ovarian and breast cancer was 35% (7,163/20,493) and 38% (14,700/38,550), respectively—significantly higher compared to the prevalence of chr13q loss of other cancer types (ranked at 12/53 and 8/53, p = 0.0206 and p = 0.0007, respectively, Permutation test). Overall, evidence obtained across the three large-scale cohorts demonstrates that breast and ovarian cancer tissues have a significantly higher prevalence of chr13q and chr17q loss than other cancer types.

### Interactions between chromosomal deletion and *gBRCA1/2* variant status

In tumors with gBRCA1/2, LOH caused by a chr17q/13q deletion suggests a carcinogenic two-hit event. However, it is also possible that chr17q/13q deletion events occur and are positively selected regardless of germline BRCA1/2 status. To investigate this, we analyzed the overlaps between gBRCA1/2 and chr17q/13q deletions. In TCGA, 2 out of 21 gBRCA1 carriers had chr17q deletions in breast cancer, and 9 out of 34 gBRCA1 carriers had chr17q deletions in ovarian cancer. Additionally, 3 out of 19 gBRCA2 carriers had chr13q deletions in breast cancer, and 12 out of 27 gBRCA2 carriers had chr17q deletions in ovarian cancer (Figure 2A). However, given this sample size, we were unable to detect any significant co-occurrence of chr17q/13q deletions with gBRCA1/2 (Supplementary Table 3). The proportion of TCGA ovarian and breast tumors with chr17q or chr13q deletions (ranging from 19% to 72%) is much higher than the percentage of gBRCA1/2 carriers—gBRCA1: 2% in breast and 8.7% in ovarian cancer; gBRCA2: 1.9% in breast and 6.6% in ovarian cancer.

**Figure 2.**
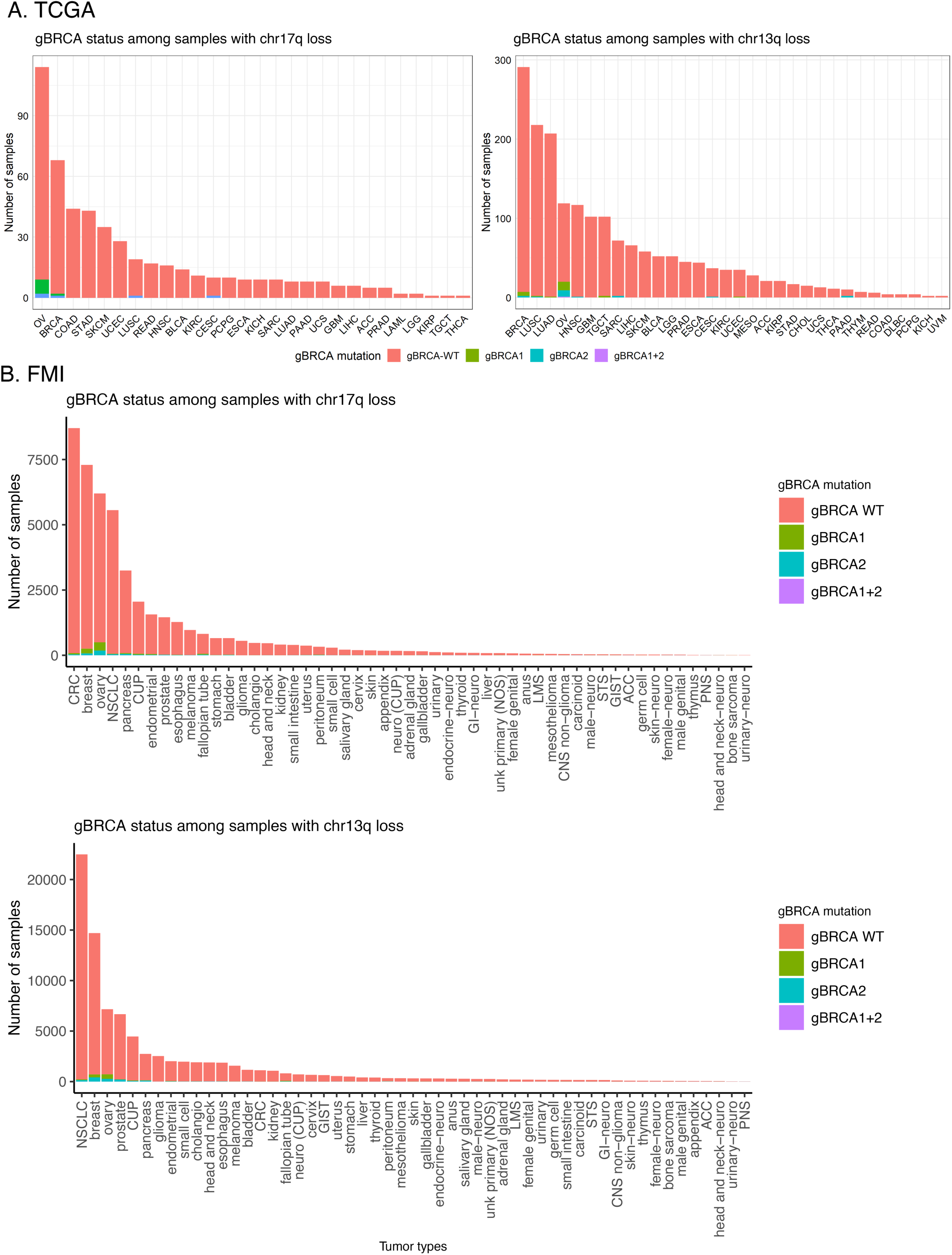
Distribution of *gBRCA1/2* status among samples with chromosome arm 17q (left panel) and 13q (right panel) deletions in the (A) TCGA and (B) FCore cohorts.

In the FCore cohort, germline *BRCA1* and *BRCA2* mutations were generally rare (**Figure 2B**, **Supplementary Tables 4-7**). However, breast and ovarian cancer tumors with chr17q or chr13q arm losses had the highest number of cases with *gBRCA1/2*. Among cases with chr17q loss, *gBRCA1/2* were most commonly seen in ovarian cancer (n = 495 cases), followed by breast cancer (n = 255 cases) (**Figure 2B**, **Supplementary Table 4**). Among cases with chr13q loss, *gBRCA1/2* were seen in several cases of ovarian (n = 715 cases) and breast cancer (n = 711 cases) (**Supplementary Table 5**). Of note, loss of chr17q significantly co-occurred with *gBRCA1* in breast, ovarian, bladder, and endometrial cancer (**Supplementary Fig. S2A**, **Supplementary Table 6**), and loss of chr13q significantly co-occurred with *gBRCA2* variants in ovarian, pancreatic, breast, and prostate cancers (**Supplementary Fig. S2B**, **Supplementary Table 7**). These patterns suggest that chromosomal arm loss may contribute to the mechanisms of biallelic inactivation of *BRCA1/2* genes. Again, the fraction of ovarian and breast cancer cases with chr17q loss (ovarian: 30.3%; breast: 18.9%) and chr13q loss (ovarian: 35%; breast: 38%) was much greater than cases with *gBRCA1* (ovarian: 4.5%; breast: 1.8%) and *gBRCA2* (ovarian: 2.5%; breast: 2.6%) in the FCore cohort. It is possible that the breast and ovarian tissue environments may be more inducive or more selective for chr17q and chr13q deletions, leading to LOH and tissue-specific cancer risk in *gBRCA1/2* carriers.

One possible positive selective advantage is that these chromosome arms contain more significantly mutated driver genes that are tumor suppressor genes (TSGs) of these cancer types, and thus their loss confers a higher selective advantage. TCGA PanCanAtlas project identified 19 TSG cancer driver genes^24^ for breast cancer, among which *BRCA1* (17q21.31), *NF1* (17q11.2), and *RB1* (13q14.2) were located on these arms. Notably, for ovarian cancer, 5 TSGs were identified as drivers by TCGA PanCanAtlas^24^. Among these five driver genes, *BRCA1*, *NF1*, *RB1*, and *CDK12* (17q12) were all located on chr17q or chr13q, except for *TP53* (17p13.1). Thus, these arm-level deletions will be sufficient to disrupt most of the known TSG driver genes in ovarian cancer.

### Chromosome-arm deletions at the cellular level and timing

To determine whether chr17q and chr13q deletions arise early in tumorigenesis or confer a strong selective advantage in breast and ovarian cells, we analyzed single-cell whole-genome sequencing (scWGS) data across multiple cancer types and pre-malignant tissues.

We examined two scWGS datasets. The first, from Funnell *et al.*^26^, comprised Triple-Negative Breast Cancer (TNBC) samples, High-Grade Serous Ovarian Cancer (HGSC) samples, and genetically engineered 184-hTERT mammary epithelial cell models. The TNBC cohort included 4 FBI (enriched in fold-back inversions), 2 HRD-Dup (enriched in small tandem duplications, including one gBRCA1 and one sBRCA1), and 1 TD (enriched in large tandem duplications) sample. The HGSC cohort included 8 FBI, 6 HRD-Dup (including one gBRCA1 and one sBRCA1), and 2 TD samples. The second dataset, from Williams *et al.*^17^, comprised luminal breast epithelial cells from gBRCA1, gBRCA2, and wild-type carriers, including samples from benign, DCIS, ALH, and LCIS tissue (**Supplementary Table 8**, **Supplementary Figure 3A**).

Most TNBC and HGSC samples had a low fraction (<2%) of cells with chr17q deleted; however, the samples that were affected showed a high percentage of cells harboring these deletions (**Figure 3A, 3C**). In the TNBC HRD-Dup sample carrying a gBRCA1 variant, 94% of cells (2321/2473) exhibited at least 25% of chr17q deleted, and 36% (896/2473) displayed over 50% deletion (**Figure 3A**). In the HGSC-gBRCA1 sample, 28% of cells (156/556) had at least 25% of chr17q deleted, with 27% (149/556) exceeding 50% deletion (**Figure 3C**). The HGSC-sBRCA1 sample showed 3% of cells (29/990) with over 25% deletion, while the HGSC-TD sample SA1047 was the most extensively affected, with 99% of cells (342/347) displaying at least 50% deletion. Among the luminal gBRCA2 samples from Williams *et al.*^17^, chr17q deletions were rare: in B2.16, only 0.5% of cells (1/189) exceeded 50% deletion, and in B2.18, no cells surpassed 50%, although 1.3% (1/79) had at least 25% deleted (**Figure 3E**).

**Figure 3.**
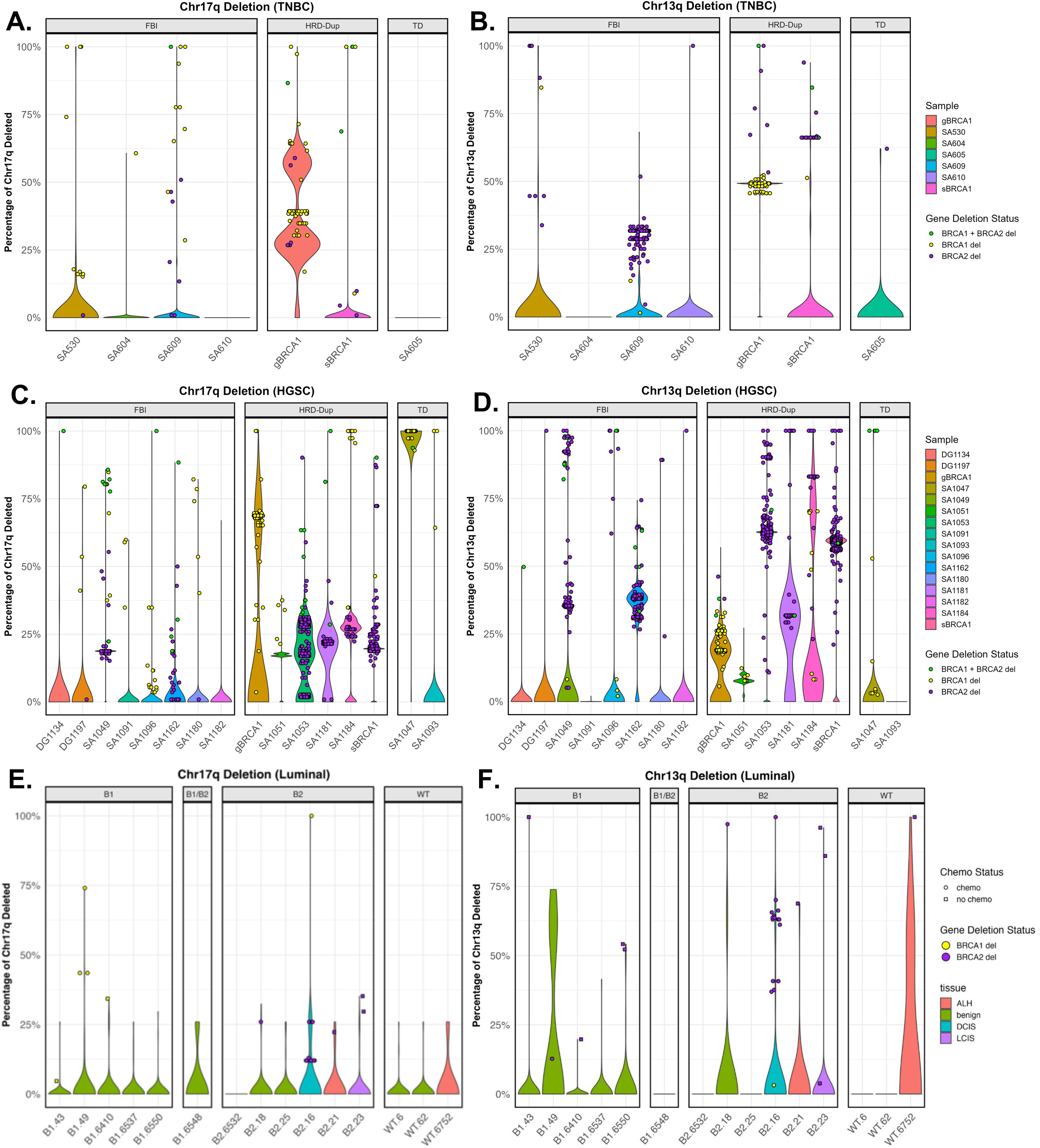
Single-cell analysis of copy number deletions on chromosome arms 17q and 13q in triple-negative breast cancer (TNBC) and high-grade serous carcinoma (HGSC) samples. Percentage of single cells harboring copy number deletions on chromosome arms (A) 17q and (B) 13q in TNBC samples based on scWGS data. Percentage of single cells harboring copy number deletions on chromosome arms (D) 17q and (C) 13q in HGSC samples based on scWGS data. Percentage of single cells harboring copy number deletions on chromosome arms (E) 17q and (F) 13q in luminal epithelial samples from scWGS data. Plots compare the percentage of CNDs between gBRCA1 tumor cells and gBRCA1 preneoplastic normal cells.

Chr13q deletions were more widespread across samples, with variable fractions of cells affected (**Figure 3B, 3D**). The TNBC-gBRCA1 sample showed the highest burden, with 94% of cells (2330/2473) exhibiting over 25% of chr13q deleted (**Figure 3B**). In the TNBC-sBRCA1 sample, 5% of cells (96/1801) had over 25% deletion, and the TNBC-FBI sample SA609 showed 3.5% (212/6033). In the HGSC cohort, the gBRCA1 sample had 9% of cells (49/556) with at least 25% deletion (**Figure 3D**), while the sBRCA1 sample was heavily affected, with 83.4% of cells (826/990) exceeding 25% and 82.9% (821/990) exceeding 50%. The HGSC-FBI sample SA409 showed 8% of cells (108/1283) with over 25% deletion. In the luminal gBRCA2 samples, B2.16 had 0.5% of cells (1/189) exceeding 50% of chr13q deleted, while in B2.18, no cells surpassed 50%, although 1.3% (1/79) had at least 25% deleted (**Figure 3F**). Overall, although the fraction of chr17q- or chr13q-deleted samples in this small scWGS dataset was lower than in the TCGA/ICGC/FCore cohorts, the affected samples showed that these deletions involved a substantial fraction of cells.

We next assessed the cancer type-specific timing of these deletions using mutational timing data from Gerstung *et al.*^25^, which ranked the relative timing of recurrent somatic events (*i.e.,* those occurring in >5% of samples) across 24 cancer types with at least ten ranked events (**Supplementary Table 9**). In ovarian adenocarcinoma, chr17 deletion ranked 1st and chr13q deletion ranked 5th, placing both among the earliest events. In breast adenocarcinoma, chr13q deletion ranked 9th, whereas chr17 deletion was not independently timed. By comparison, among the remaining 22 cancer types, only chromophobe renal cell carcinoma ranked chr17 deletion (3rd). For chr13q deletion, chronic lymphocytic leukemia (3rd), chromophobe renal cell carcinoma (4th), and osteosarcoma (4th) ranked it as an early event; all other cancer types placed it late (≥17) or left it unranked. These results demonstrate that chr17 and chr13q deletions are among the earliest somatic events in breast and ovarian cancers relative to other malignancies.

Consistent with the first-hit hypothesis, these deletions are most apparent at the earliest stages of breast cancer. In the scWGS data from Williams *et al.*^17^, DCIS and benign luminal epithelial samples from gBRCA2 carriers (B2.16 and B2.18) showed phylogenetic trees with small, localized clades of cells harboring chr13q and/or chr17q deletions, while the majority of cells remained CNV-neutral. The corresponding genome-wide copy-number heatmaps displayed relatively sparse overall alterations (**Fig. 4 A–D**).

**Figure 4.**
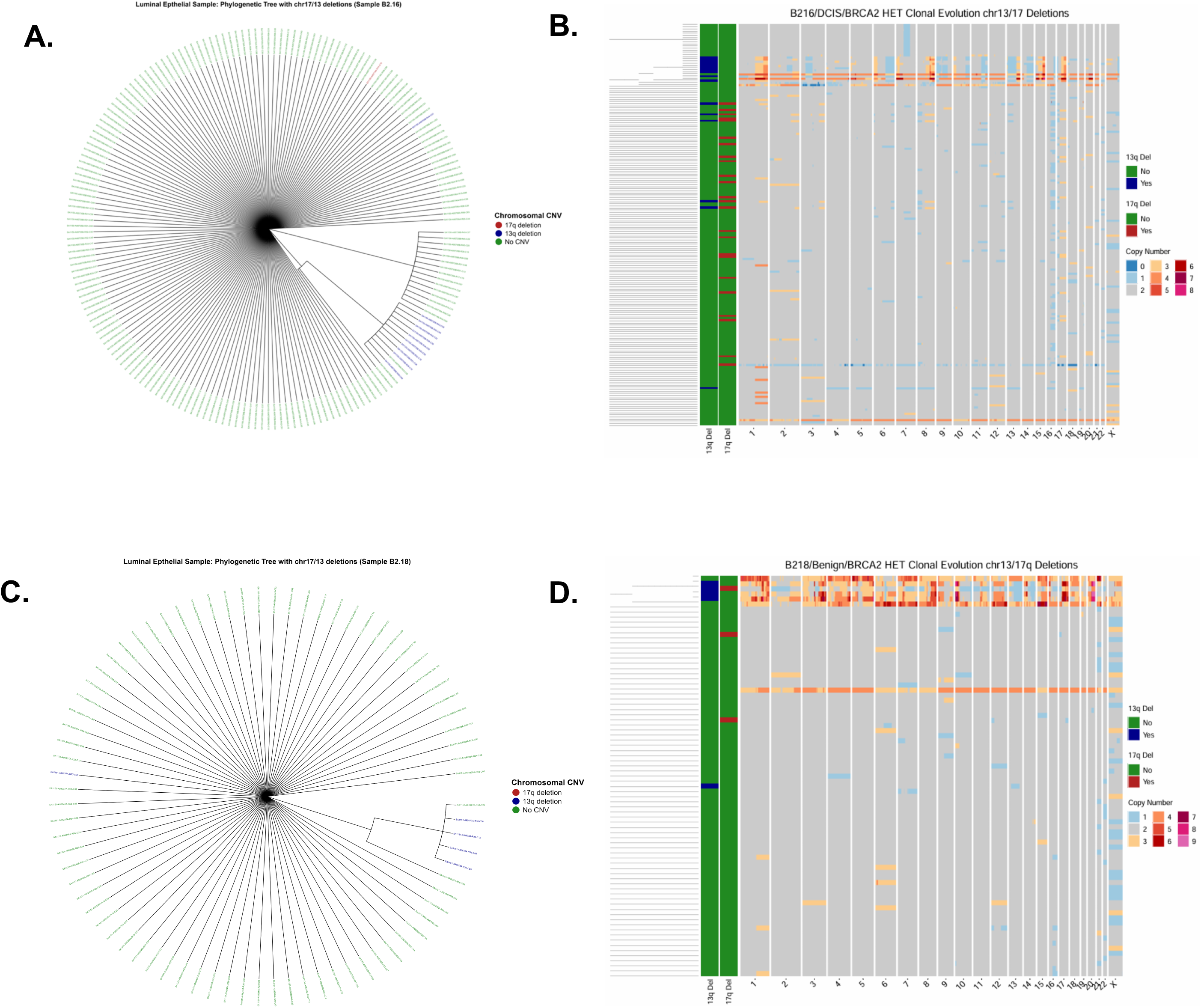
Clonal Architecture of Chromosomal Deletions in Pre-malignant Breast Tissue. Phylogenetic trees of luminal epithelial cells from a BRCA2/HET carrier with DCIS (Sample B216, **A**) and benign tissue (Sample B218, **C**). Cells are clustered based on copy number profiles **from scWGS data**. Tip colors indicate the presence of specific structural variations: chr17q deletion (red), chr13q deletion (blue), or no detected CNV (green). The trees show small, distinct clades with 17q and 13q deletions, suggesting these events occur in localized sub-clones prior to full malignancy. **(B, D).** Genome-wide copy number heatmaps for the corresponding samples. The left-hand sidebar indicates cells with 13q (top bar) and 17q (bottom bar) deletions. While most of the genome remains diploid (gray), early deletions in chromosomes 13 and 17 are visible as focal losses against a relatively stable genomic background.

We further assessed the phylogenetic distribution of chr17q and chr13q deletions in the Funnell *et al.*^26^ scWGS data, restricting the analysis to cells with at least 50% q-arm loss. Clonality was evaluated using the Fritz and Purvis D statistic. For chr13q, deletions were phylogenetically dispersed in both TNBC and HGSC, consistent with late, independent acquisition. In the gBRCA1 TNBC sample SA501, chr13q deletions showed no phylogenetic structure (D = 0.98; p₁ = 0.568, p₀ = 0.00; **Fig. 5A**), and in the gBRCA1 HGSC sample SA1052, deletions were similarly dispersed (D = 3.73; p₁ = 0.9, p₀ = 0.09; **Fig. 5C**). In contrast, chr17q deletions formed well-defined clusters on the phylogenetic trees. In SA501, deletions occupied a distinct branch (D = −0.30; p₁ = 0.0, p₀ = 0.9), indicating significant clustering consistent with Brownian trait evolution. SA1052 showed similar clonality (D = 0.26; p₁ = 0.0, p₀ = 0.06). Together, these results suggest that chr17q deletions are early clonal events in germline tumors, whereas chr13q deletions arise later and independently (**Fig. 5A–D**).

**Figure 5.**
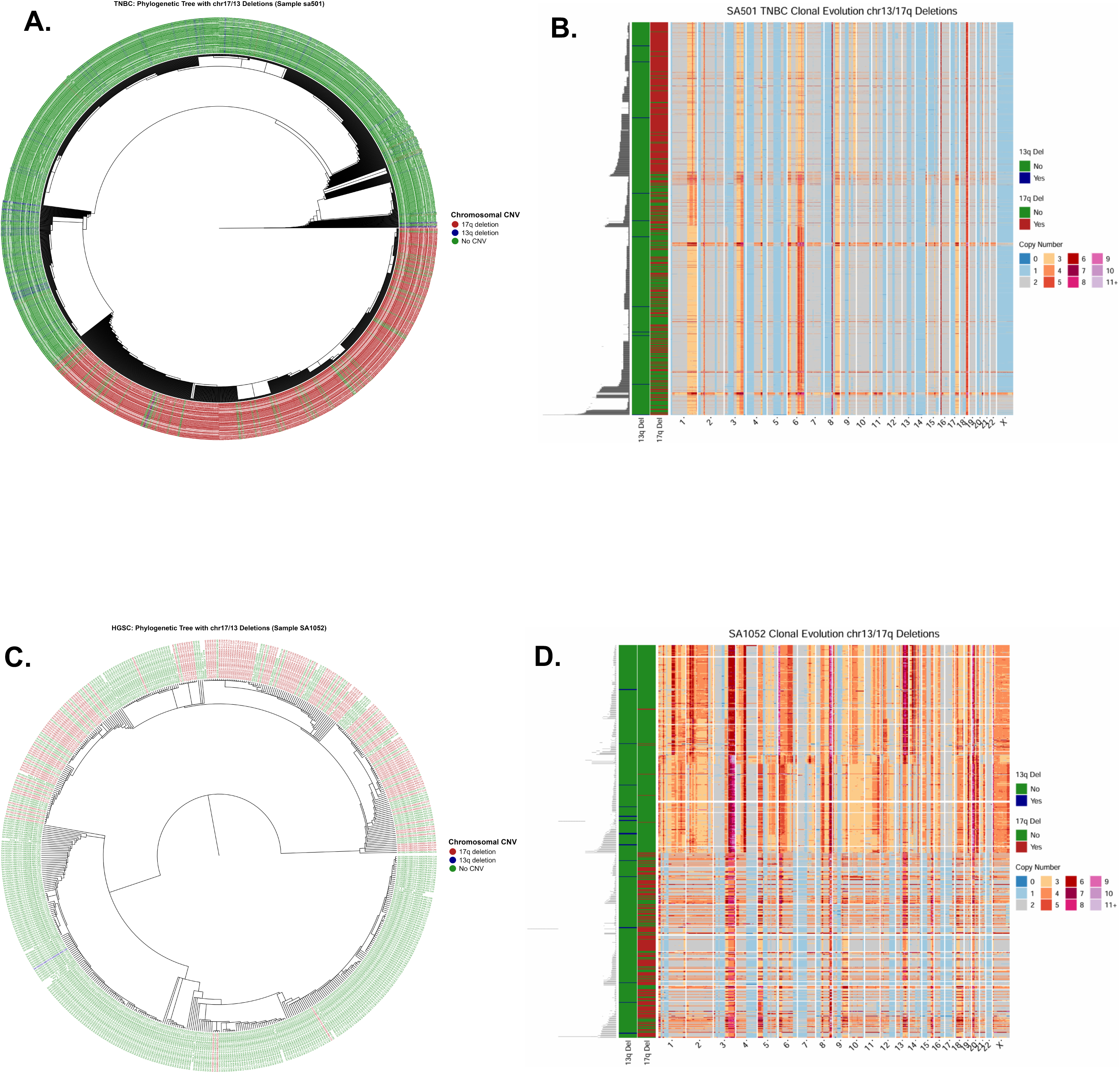
Clonal Expansion and Evolutionary Trajectories in TNBC and HGSC. **(A, C)** Circular phylogenetic trees for Triple-Negative Breast Cancer (Sample sa501, **A**) and High-Grade Serous Carcinoma (Sample SA1052, **C**). In contrast to pre-neoplastic samples, large ancestral branches are marked by the chr17q deletion (red), indicating a massive clonal sweep where cells carrying this deletion have dominated the tumor population. **(B, D).** Genome-wide copy number heatmaps demonstrating high levels of genomic instability. In both samples, the foundational 17q loss (red sidebar) is associated with widespread saneuploidy, including complex gains (orange/red) and losses (blue) across multiple chromosomes.

To assess clonality in germline BRCA1/BRCA2 contexts, we examined non-invasive mammary epithelial samples: a BRCA2⁺/⁻ 184-hTERT model (SA1188) from Funnell *et al.*^26^, and a DCIS gBRCA2 sample (B2.16) and a benign gBRCA2 sample (B2.18) from Williams *et al.*^17^. In SA1188, chr17q deletions were rare (11/556 cells) and showed limited phylogenetic structure (D = −1.34; p₁ = 0.0) while remaining consistent with Brownian evolution (p₀ = 0.91). Chr13q deletions in this sample (10/556 cells) showed no evidence of non-random distribution (D = 0.95; p₁ = 0.11) and were inconsistent with Brownian inheritance (p₀ = 0.02).

In the luminal samples, chr17q deletions were extremely sparse: B2.16 had only 1/189 deleted cells (D = 2.14; p₁ = 0.06, p₀ = 0.69), and B2.18 had 1/79 (D = −4.79; p₁ = 0.0, p₀ = 0.91), with significance in the latter likely attributable to trait sparsity rather than true clonality. In contrast, chr13q deletions in B2.16 demonstrated strong phylogenetic clustering (D = −4.79; p₁ = 0.0, p₀ = 0.91), consistent with early clonal acquisition, whereas B2.18 showed no clustering (D = −6.86; p₁ = 0.96, p₀ = 0.19), with the extreme D values again likely reflecting sparsity. Overall, chr17q deletions were rare in luminal samples, whereas chr13q deletions showed clonal structure only in B2.16.

In established malignancies such as TNBC (SA501) and HGSC (SA1052), the same deletions are found within dominant CNV-defined lineages. Deletion-positive cells occupy large fractions of the phylogeny, consistent with clonal sweeps in which a founding cell carrying these mutations gained a growth advantage and expanded to dominate the tumor population. Concurrently, the genomic landscape becomes far more complex: copy-number heatmaps display dense patterns of gains and losses across many chromosomes, reflecting widespread chromosomal instability and ongoing subclonal diversification (**Fig. 5A–D**). Together, these phylogenomic analyses link early, localized deletion events in pre-malignant tissue to later malignant states in which those events are incorporated into dominant lineages and accompanied by extensive genome-wide copy-number evolution.

## DISCUSSION

Here, we establish correlations between chromosomal deletions that encompass the BRCA1/2 gene loci and the cancer types with the highest risk for gBRCA1/2 variant carriers, using three independent datasets comprising 340,824 cancer cases. These findings suggest chr17q/13q deletions, observed in higher prevalence among breast and ovarian cancer, may serve as a “second hit” that may explain the tissue-specific cancer risk associated with gBRCA1/2. Notably, the fraction of tumors with chr17q or chr13q deletions are a magnitude greater than the fraction of gBRCA1/2 pathogenic variant carriers in breast cancer or ovarian cancer cases. It is possible that the breast and ovarian cellular or tissue environment may be more inducive or confer higher selective advantage for cells with chr17q and chr13q deletions, which lead to LOH and potential development of breast and ovarian tumors in gBRCA1/2 variant carriers.

Our single-cell analyses provide direct cellular-level support for these bulk-cohort observations. Using scWGS data from Funnell *et al.*^26^ and Williams *et al.*^17^, we showed that in affected TNBC and HGSC samples, chr17q and chr13q deletions are present in a substantial fraction of cells, consistent with clonal or near-clonal events rather than sporadic aberrations. Phylogenetic analysis using the Fritz and Purvis D statistic further demonstrated that chr17q deletions form well-defined clonal clusters in established tumors such as SA501 (TNBC) and SA1052 (HGSC), consistent with early acquisition and subsequent clonal expansion. In contrast, chr13q deletions were phylogenetically dispersed, suggesting they arise later and independently across multiple lineages (**Fig. 5**). These patterns are concordant with mutational timing data from Gerstung *et al.*^25^, which ranked chr17 deletion among the earliest somatic events in ovarian adenocarcinoma and chr13q deletion as an early event in both breast and ovarian cancers.

Importantly, our analysis of pre-malignant tissues further supports the early origin of these deletions. In DCIS and benign luminal epithelial samples from gBRCA2 carriers (B2.16 and B2.18), scWGS-derived phylogenetic trees revealed small, localized clades of cells harboring chr13q and/or chr17q deletions against an otherwise CNV-neutral background (**Fig. 4**). Chr13q deletions in B2.16 showed strong phylogenetic clustering consistent with early clonal acquisition, providing evidence that these events can precede the transition to invasive malignancy. By contrast, the established TNBC and HGSC tumors displayed dense genome-wide copy-number alterations alongside these deletions (**Fig. 5**), illustrating how early, localized deletion events become incorporated into dominant lineages accompanied by extensive chromosomal instability. Together, these observations are consistent with a model in which chr17q and chr13q deletions serve as founding events that facilitate subsequent genomic diversification.

It remains to be further examined whether these deletions also arise in normal tissues. A recent study^29^ inferred somatic copy number alterations (sCNAs) in 1,708 TCGA normal-appearing adjacent-to-tumor (NAT) tissue samples, detecting copy number gain, loss, and copy-neutral loss of heterozygosity (cn-LOH), *i.e.*, the loss of one allele concurrent with gain of the other allele. Based on this bulk-tissue analysis of sCNAs, we queried copy number loss or cn-LOH in normal adjacent tissues (NATs). Only 3 NAT samples showed CN loss on chr17q (2 ovarian cancer and 1 ESCA), and while no NAT samples showed chr13q loss, two NAT samples that showed cn-LOH on chr13q were from patients with ovarian cancer and ESCA. We note that the detection limit for this bulk tissue analysis was estimated to be ∼5–10% of cells harboring somatic DNAs; high-resolution analyses of tissues across the trajectory of disease development are required to more precisely identify the origin, timing, and possible selection of these aneuploidy events. Our scWGS findings in pre-malignant samples reinforce this point: even in DCIS and benign tissues from gBRCA2 carriers, only rare cells harbored chr17q deletions, underscoring that these events may be present at frequencies below the detection threshold of bulk approaches.

Beyond chr17q and chr13q, abnormalities of other chromosomes have also been reported in breast and ovarian cancer. A recent study of cancer/non-cancer breast tissues identified derivative chromosome der(1;16) as a key clonal event. This genomic alteration includes the entire long arm of chromosome 1 (1q) and a truncated short arm of chromosome 16 (16p), acquired during early puberty to late adolescence in breast cancer patients^30^. Analyzing published data from a recent study^31^ combining unbiased chromosome-arm genetic screens and *in vitro* evolution, chr13q loss was substantially more selected for in normal diploid human telomerase reverse transcriptase (hTERT)-immortalized breast mammary cells (HMECs, loss-gain frequency=25%) compared to renal epithelial cells (RPTECs, loss-gain frequency=0). While chr17q loss was not favored in either type of cells, chr17p loss also shows the same pattern (HMECs loss-gain freq=40.3%, RPTECs loss-gain freq=0). This data demonstrates the selective advantage of chr13q loss and chr17p loss (which may co-occur with chr17q loss through chr17 loss) in breast versus renal cells. Thus, even though renal cancer showed high prevalence of these events in our TCGA/ICGC/FCore cohort analyses, breast mammary cells may provide a specific environment where selection for these events is more favorable. Consistent with this, our scWGS analysis of the BRCA2⁺/⁻ 184-hTERT model (SA1188) from Funnell *et al.*^26^ showed that chr17q and chr13q deletions were detectable in this non-malignant immortalized mammary epithelial cell line, albeit at low frequencies, further supporting the notion that the breast epithelial environment permits the emergence of these alterations.

Our findings have potential clinical implications in the context of cancer risk assessment and targeted therapies. The high prevalence of chr17q and chr13q deletions in breast and ovarian cancers suggests that these chromosomal alterations could serve as biomarkers for early detection or risk stratification. The single-cell evidence that these deletions can be detected in pre-malignant tissues and that they exhibit clonal structure raises the possibility that monitoring for such events in at-risk individuals—for example, through high-sensitivity assays of breast tissue biopsies from gBRCA1/2 carriers—could identify individuals at elevated risk of progression. Additionally, understanding the mechanisms by which these deletions contribute to tumorigenesis could inform the development of targeted therapies aimed at exploiting vulnerabilities in tumors with these specific chromosomal alterations. Moreover, our results underscore the importance of considering chromosomal instability and its interplay with tissue-specific factors in tumourigenesis.

While our results provide a plausible mechanistic link between breast cancer LOH and cancer risk, there were several limitations. While genomic cohorts can help identify associations, they do not suggest causality. It remains possible that the elevated prevalences of chr17q and chr13q deletions in ovarian and breast cancers might be a consequence rather than a cause, although evidence from mutational timing and our phylogenetic analyses suggest these events occur early. Further, other BRCA1/2-associated cancer types, including PRAD and PAAD, show less consistent patterns of higher chr17q/13q deletion prevalences across cohorts (**Figure 1**), albeit the relative risks for these cancer types are substantially lower^8,9^. The single-cell datasets analyzed here, while informative, were limited in sample size; larger cohorts with matched normal, pre-malignant, and tumor tissues will be needed to more precisely characterize the timing and clonal dynamics of these deletions. Future studies may consider studying aneuploidy in normal and pre-cancer tissues at high-resolution and in larger sample sizes, or utilize experimental methods to introduce arm-level loss and determine causality^31^.

In conclusion, this study sheds light on the prevalence and significance of chr17q and chr13q deletions in breast and ovarian cancers, highlighting their potential role in tissue-specific cancer risk associated with germline *BRCA1/2* mutations. By integrating bulk-cohort prevalence data with single-cell phylogenetic evidence, we show that these deletions are not only highly recurrent but also exhibit patterns of early clonal acquisition and expansion—particularly for chr17q—consistent with a founding role in tumorigenesis. Our findings contribute to a new hypothesis that may explain tissue-specific risk associated with gBRCA1/2 predisposition. Further studies are needed to elucidate the functional consequences of these deletions and their potential as biomarkers or therapeutic targets in cancer treatment.

## METHODS

### Genomic datasets TCGA

TCGA somatic ploidy and CNV data: We collected arm-level and gene-level CNV calls of 10,713 cases as well as corresponding whole-genome doubling profiles from TCGA PanCanAtlas Aneuploidity Working Group^32^. To obtain this CNV data, the likeliest ploidy was determined by the ABSOLUTE method in the genome for each tumor. Meanwhile, the CNV data were generated from results of Affymetrix SNP 6.0 arrays. To obtain arm-level CNV events, the absolute copy numbers of each segment in the genome was calculated using the ABSOLUTE algorithm. Then, a Gaussian Mixture Model implemented with the Python package SciKit-Learn was used to combine arm/chromosome data and determine alterations. In our research, deletions (-1) of chr13q and chr17q were used to calculate prevalences. For gene-level CNV events, thresholded deletion values generated by GISTIC2^34^ calls were used for BRCA1/BRCA2 gene deletion status, where both deep deletions (-2) and shallow deletions (-1) at the regions of these 2 genes were considered as deletions for further calculation. Based on this, we filtered for primary tumor samples from unique cases, yielding a final of 10,522 tumors for analyses.

TCGA germline variant data: We obtained *BRCA1/2* pathogenic/likely-pathogenic germline variants that were identified in TCGA cases from the PanCanAtlas Germline Working Group ^23^. Variants were downloaded upon dbGaP approval from GDC.

### ICGC PCAWG

ICGC somatic ploidy and CNV data: We obtained arm-level CNV events of 2,507 patients and corresponding WGD profiles from ICGC PCAWG project (https://dcc.icgc.org/releases/PCAWG). The detailed methods used for ICGC somatic copy number data were described in a previous study^35^. Briefly, the ploidy profile of each sample was generated by ascatNgs. Then, consensus copy number profiles were generated from the combination of outputs of 6 copy-number alteration algorithms using a multi-tiered approach. In our analysis, deletions (copy number values less than or equal to -1) of chr13q and chr17q in different samples were collected to calculate prevalences. In the final ICGC samples, we retained only one primary tumor per patient that had no overlapping samples in TCGA to ensure independence in these two analyses.

### FoundationCore® (FCore) database (Foundation Medicine comprehensive genomic profiling cohort)

A total of 328,573 patients with solid tumors who received comprehensive genomic profiling (CGP) using FoundationOne^®^/FoundationOne^®^CDx assays, as part of routine clinical care, through March 2023 were included in this study. Approval for this study, including a waiver of informed consent and a Health Insurance Portability and Accountability Act waiver of authorization, was obtained from the WCG Institutional Review Board (Protocol No. 20152817).

DNA was extracted from Formalin-fixed, paraffin embedded (FFPE) tumor sections. 50-1000 ng of DNA underwent whole-genome shotgun library construction and hybridization-based capture of all coding exons from 309 cancer-related genes, one promoter region, one non-coding (ncRNA), and select intronic regions from 34 commonly rearranged genes, 21 of which also include the coding exons, as described previously^36,37^. Sequence data were analyzed for base substitutions, short insertions/deletions, copy number changes and gene rearrangements. Specifically, for the assessment of pathogenic germline *BRCA1/2* alterations in this study, the prediction from the somatic-germline-zygosity (SGZ) algorithm^38^, as well as a consensus germline prediction overall or in any ancestry group based on >600,000 samples sequenced at FMI was utilized^39^.

For detecting chr13q and chr17q arm loss events, a copy-number modeling algorithm described previously was used^40,41^. Briefly, the algorithm utilized coverage data of the targeted regions of the genome, normalized to a process-matched control, to model the copy number of each segment. Along with the normalized intensities, the minor allele prevalences of the SNPs distributed across each segment were used to identify regions under aneuploidy (gain, loss). If >50% of the arm exhibited aneuploidy, it was considered a chromosome arm event.

### Permutation test to assess the significant prevalence of arm-level loss

In TCGA and ICGC, we conducted a permutation test to assess the prevalence of chromosome arm deletions, particularly chr13q and chr17q, across cancer types. The permutation test was designed to determine the significance of observed deletion rates within each cancer type by comparing them against a null distribution generated through random shuffling of the data. For each cancer type and chromosome arm, we calculated the observed proportion of deletions and generated a null distribution through 10,000 permutations. This involved shuffling the combined pool of deletion data from all samples in the full and non-WGD datasets, thereby simulating a distribution of deletion proportions under the null hypothesis. The p-values were derived by comparing the observed deletion proportions to the permutation-generated null distribution. To account for multiple testing, we applied the Benjamini-Hochberg procedure to adjust the p-values.

In the FCore cohort, to test the ranks of prevalences of chr13q or chr17q deletions in different cancer types, we performed permutation tests. Firstly, we shuffled the cancer-type labels for each patient and calculated the prevalence of indicated chromosomal-arm deletions for each cancer type. Then, we obtained the ranks of each cancer type based on the prevalence of chr13q or chr17q deletions. By repeating the above steps for 10,000 iterations, we generated 10,000 random ranks. Finally, the number of times that the prevalences of indicated chromosomal arm (chr13q or 17q) deletions in breast cancer or ovarian cancer ranked higher than or equal to their real ranks was counted, and then divided by 10,000 to obtain the corresponding significance level, i.e., P values.

### Mutational Timing Analysis of Chromosomal Arm Deletions

We utilized mutational timing estimates from Gerstung *et al*^25^. to analyze the sequence of chr13q and 17 deletions in breast and ovarian cancer. Specifically, Gerstung et al. focused on recurrent events occurring in more than 5% of samples across cancer types. To estimate the relative timing of these mutations, they constructed a discrete multinomial distribution based on the pairwise probabilities of one mutation occurring before another. Using a league model approach, they simulated a series of “games” by drawing from this multinomial distribution. Points were assigned based on “wins” (when mutation A was determined to occur before mutation B) or “ties” (when the order was unknown). The final scores from these simulations, repeated 1,000 times, were ranked to generate relative timing estimates for the mutations. Here, we extract these results and evaluate the timing of chromosome 13q and 17 deletions in each cancer type for which at least 10 events were timed.

### Single-cell whole-genome sequencing (scWGS) Dataset and Analysis

We utilized the two single-cell Whole Genome Sequencing (scWGS) datasets from Funnel et al^26^. which included Triple Negative Breast Cancer (TNBC), High-Grade Serous Ovarian Cancer (HGSC) and genetically engineered Mammary Epithelial Cells (184-hTERT) datasets. The TNBC dataset provided 7 samples, with 4 containing fold-back inversions (FBIs), 2 BRCA1 samples with homologous recombination deficiencies (HRD-Dup) and 1 sample enriched in large tandem duplications (TD). In addition, we analyzed 16 HGSC samples, including 8 samples with FBIs, 8 exhibiting HRD-Dups (including 2 BRCA1 samples), and 2 with TDs. Lastly, the 184-hTERT Mammary Epithelial Cell datasets provided cells with genetically engineered BRCA1, BRCA2, and TP53 deletions via CRISPR–Cas9 editing of 184-hTERT L9 cell lines.

Additionally, allele-specific copy number profiles of 15 luminal breast epithelial samples from Williams et al^27^. were included. These samples consist of benign tissues from germline BRCA1 carriers (B1-43, B1-49, B1-6410, B1-6537, B1-6550), BRCA2 carriers (B2-18, B2-25, B2-6532), and wild-type individuals (WT-6, WT-62). Precancerous lesions included one ductal carcinoma in situ (B2-16), one lobular carcinoma in situ (B2-23), and two atypical lobular hyperplasias (B2-21, WT-6752). Lastly, one benign sample (B1-6548) possessed germline mutations in both BRCA1 and BRCA2.

With the scWGS datasets provided, copy number (CN) states were inferred by the R program, HMMcopy. The number of megabases affected by copy number deletion (CND) was calculated from CN states 1 and 2, whereas states 4-11 were considered copy number amplifications or gains. We next calculated the percentage of chr13q or chr17q deletion across all cells in each sample using R.

### Statistical analysis

Descriptive statistics were performed and illustrated by using R (version 4.0.3) and python (version 3.9). In each cohort, co-occurrence and mutual exclusivity analysis of chr17q/13q arm loss with pathogenic germline variants in BRCA1/2 was performed using a two-sided Fisher’s exact test. P-values were adjusted for multiple hypothesis correction using the Benjamini–Hochberg FDR procedure. Statistical significance was set at an FDR-corrected p ≤ 0.05.

Cell phylogenies were inferred through the Sitkatree program from the CNV data of scWGS samples of interest. Each CNV was encoded as a binary trait for deletion (presence/absence) and was mapped to its corresponding cell tip. Cells were labeled with chr17q/chr13q deletions only if at least 50% of the q arm was deleted. The phylogenetic signal of the binary CNV data was quantified using the Fritz and Purvis D statistic from the R package, caper. The D statistic compares the observed distribution of a binary trait across a phylogeny with both the random distribution of a trait (D = 1) and the Brownian motion model (D = 0). Brownian motion is a model for gradual inheritance of a trait along branches, so that traits that arise early are passed to descendant cells. Caper also assessed the significance of D by testing it against the random distribution null hypothesis (P₁) and a Brownian motion null hypothesis (P₀), where a significant P₁ indicates clustering of CNVs along the phylogeny and a non-significant P₀ indicates consistency with early clonal acquisition. Statistics, computation, and plotting were carried out using Python 2.7.18 (Python Software Foundation) and R 4.2.3 (R Foundation for Statistical Computing).

Phylogenetic trees were generated with **Sitka** with default parameters. Sitka performs Bayesian phylogenetic inference from site dependencies induced by rearrangements and using scWGS copy-number data, leveraging the Sitka transformation to reduce CN site dependencies, and uses scalable MCMC to sample from the posterior over tree topologies <u>(Sitkatree)</u>

Heatmaps were generated using ComplexHeatmap in R; rows and columns were hierarchically clustered and visualized with the default settings

### Aneuploidy Analysis

Using each cell’s baseline copy number as the reference, we summarized genome-wide CNV burden by measuring the length-weighted fraction of the genome with copy number at least one below baseline (loss) or at least one above baseline (gain), and by quantifying the typical deviation size using the length-weighted median. Here, the baseline copy number represents the cell’s overall ploidy—estimated as the segment-length–weighted median CN state across the genome—so it is the CN level covering most of the genome in that cell (e.g., ∼2 in diploid cells, ∼4 in WGD/endopolyploidy cells), and gains/losses are defined relative to this cell-specific reference.

In addition to these genome-wide summaries, we implemented a chromosome-specific deletion-calling procedure to capture recurrent large-scale events. For a given target chromosome (chr13 or 17), we computed, in each cell, the fraction of that chromosome’s segmented length in a deletion-like state (by default, CN <= 1) and labeled the chromosome as deleted if this fraction exceeded a user-defined threshold (default, i.e., at least half of the chromosome).

We fit a pooled linear regression across all cells to assess whether genome-wide CNV burden (altered fraction) differs by chromosome 13 and chromosome 17 deletion status, while controlling for WGD. To obtain inference robust to heteroskedasticity, we recomputed the coefficient tests using HC3-robust standard errors and p-values. The regression coefficients quantified the adjusted effect size of each event on the fraction of the genome altered (in absolute fraction units), conditional on the other covariates in the model.

## Supplementary Figures

**Supplementary Figure 1.**
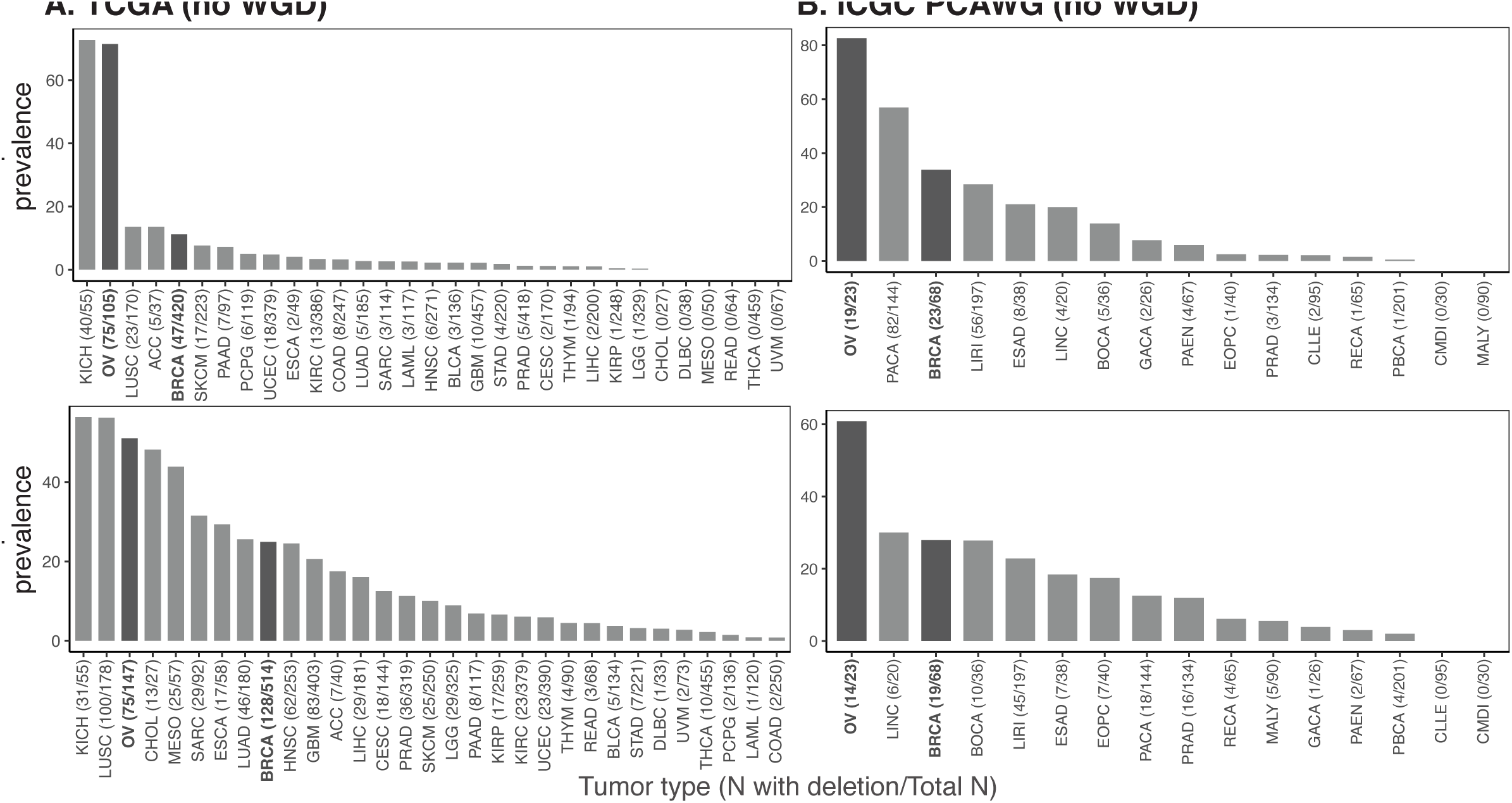
The prevalence of chr17q and chr13q deletions across tumor types in non-WGD samples of the (A) TCGA and (B) ICGC PCAWG cohorts. The x-axis lists the tumor types along with the number of samples with deletions out of the total number of samples analyzed, e.g., “CANCER (#N affected/#Total N)”. The y-axis indicates the percentage of samples with deletions for each tumor type. Tumor types with a higher prevalence of deletions are shown on the left, decreasing towards the right.

**Supplementary Figure 2.**
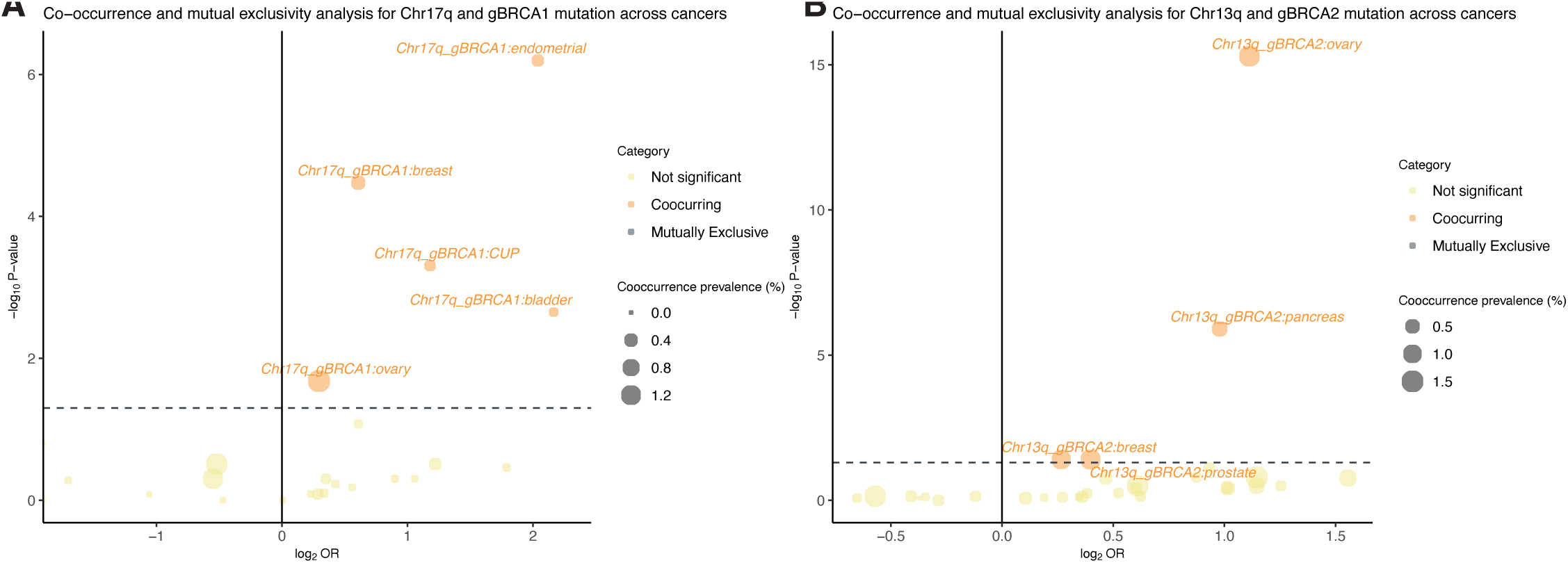
Co-occurrence and mutual-exclusivity analyses using FCore cohort to evaluate the statistical associations between (A) chr17q deletion and gBRCA1 and (B) chr13q deletion and gBRCA2.

**Supplementary Figure 3.**
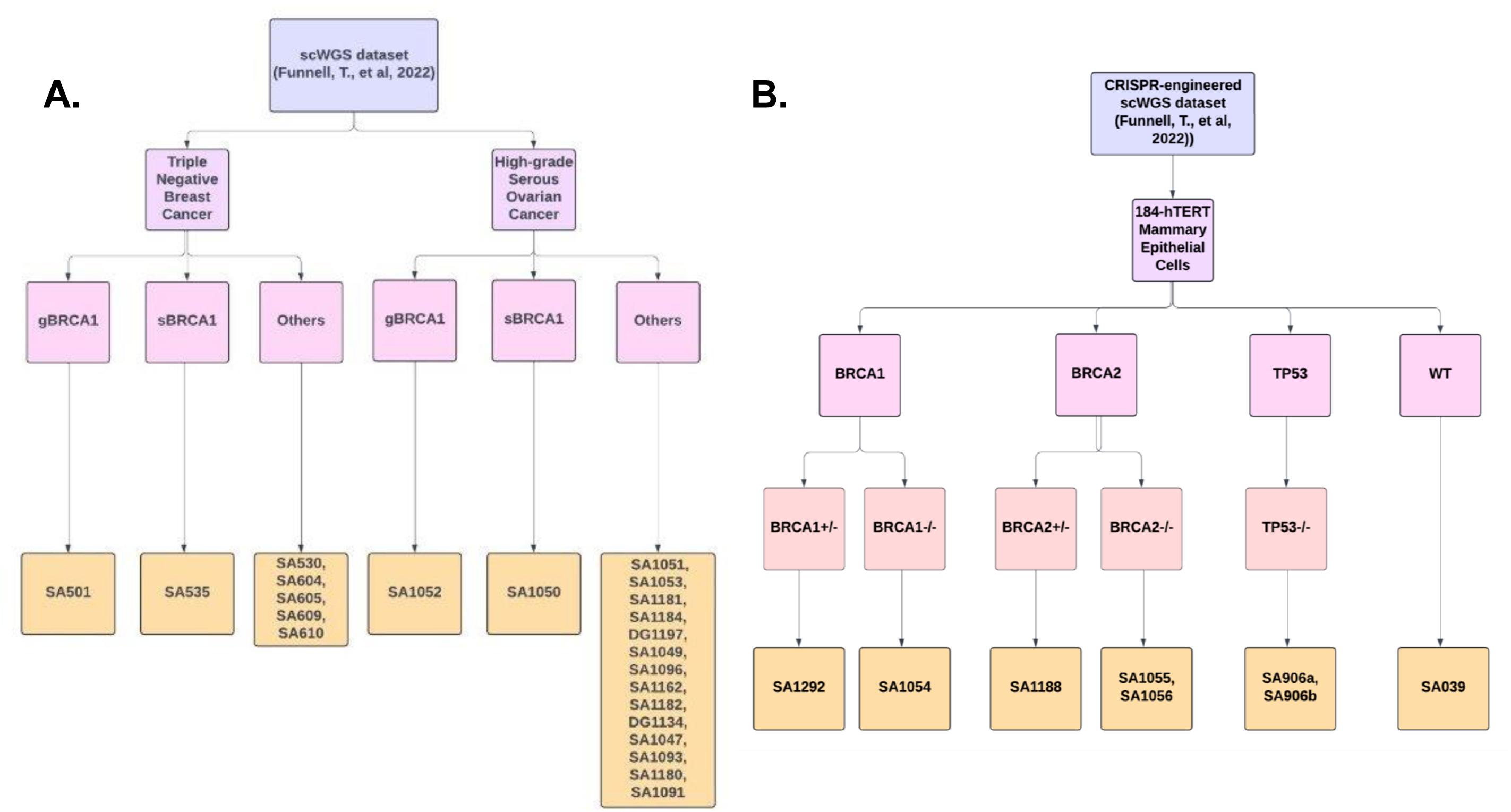
Single cell analysis of copy number deletions of chromosome arms 17q and 13q in gBRCA1/2 vs non-carrier tumor samples and genetically engineered cells. (A) Summary flowchart of single-cell datasets and samples used in analyses. The left panel displays data from Funnell, T. et al. (2022), including scWGS-analyzed both TNBC and HGSC cells possessing gBRCA1, sBRCA1 and WT BRCA1 variants. The right panel shows data from William et al. (2024). Percentage of inferred single-cell copy number deletions on chromosome arms (B) 17q and (C) 13q in TNBC samples inferred from scRNA-seq data. Plots compare the percentage of CNDs between gBRCA1 tumor cells and non-carrier tumor cells.

## DATA AVAILABILITY

Data for the TCGA PanCanAtlas somatic mutations and CNV calls are available at NCI genome data commons (GDC): https://gdc.cancer.gov/about-data/publications/pancanatlas. The controlled-access TCGA germline variants are available at https://gdc.cancer.gov/about-data/publications/PanCanAtlas-Germline-AWG for investigators with dbGaP approval. In addition, ICGC PCAWG data is available via ICGC data portal: https://dcc.icgc.org/pcawg/.

The sequencing data from Foundation Medicine Inc. utilized for this study are derived from clinical samples. The relevant data supporting the findings of this study are presented within the main article and the supplementary files. Due to the Health Insurance Portability and Accountability Act requirements, we are not authorized to share underlying sequence data or individualized patient genomic data that contain potentially identifying or sensitive patient information. Foundation Medicine is committed to collaborative data analysis, with well-established and widely used mechanisms that enable investigators to query our core genomic database of >700,000 de-identified sequenced cancers to obtain aggregated datasets. For more information and mechanisms of access, please contact the corresponding author(s) or the Foundation Medicine, Inc. Data Governance Council at.

## CODE AVAILABILITY

Code for conducting and reproducing this study can be found at: https://github.com/Huang-lab/TissueSpecificCancerRisk

## ACKNOWLEDGEMENTS

The authors wish to acknowledge TCGA, ICGC, FMI studies and their participating patients and families that generously contributed the data. The authors thank all members of the Huang lab for constructive discussion. Large language models (LLM) may have been used in the initial drafts of coding and writing of this work. All final codes and texts have been extensively edited and verified by the authors.

## CONTRIBUTIONS

K.H. conceived the research and designed the analyses. X.W., S. Sisoudiya, M.B., Y.G., K.H., and T.J. conducted the bioinformatics analyses. S. Sivakumar, and K.H. supervised the study. K.H., S. Sisoudiya, T.J., S. Sivakumar, and X.W. wrote the manuscript. All authors read, edited, and approved the manuscript.

